# Localization of neutral evolution: selection for mutational robustness and the maximal entropy random walk

**DOI:** 10.1101/2020.01.28.922831

**Authors:** Matteo Smerlak

## Abstract

If many mutations confer no immediate selective advantage, they can pave the way for the discovery of fitter phenotypes and their subsequent positive selection. Understanding the reach of neutral evolution is therefore a key problem linking diversity, robustness and evolvability at the molecular scale. While this process is usually described as a random walk in sequence space with clock-like regularity, new effects can arise in large microbial or viral populations where new mutants arise before old ones can fix. Here I show that the clonal interference of neutral variants shuts off the access to neutral ridges and thus induces localization within the robust cores of neutral networks. As a result, larger populations can be less effective at exploring sequence space than smaller ones—a counterintuitive limitation to evolvability which invalidates analogies between evolution and percolation. I illustrate these effects by revisiting Maynard Smith’s word-game model of protein evolution. Interestingly, the phenomenon of neutral interference connects evolutionary dynamics to a Markov process known in network science as the maximal-entropy random walk; its special properties imply that, when many neutral variants interfere in a population, evolution chooses mutational paths—not individual mutations—uniformly at random.

## Introduction

Kimura famously championed the view that a large part of all evolutionary change in genomes confer no selective advantage, *i.e.* molecular evolution is largely *neutral* (Kimura, 1983). Initially based on high observed substitution rates (Kimura, 1968; King and Jukes, 1969), this hypothesis was supported by the later discovery of extended neutral networks—sets of sequences connected by one-point mutations with equivalent phenotype or function (Smith, 1970)—in many molecular genotype-to-phenotype maps, e.g. in RNA secondary structure (Fontana et al., 1993), protein structure (Babajide et al., 1997) or transcriptional regulation networks (Ciliberti et al., 2007). While the exact rate of adaptive vs. neutral evolution remains under investigation (Eyre-Walker, 2006), neutralism is now widely understood as a central aspect of evolutionary dynamics (Nei et al., 2010). Besides ensuring a high level of mutational robustness (van Nimwegen et al., 1999), neutral evolution can enhance evolvability by providing access to novel—and possibly fitter—phenotypes (Wagner, 2008). In this way, the variation generated by neutral evolution enables the “arrival of the fittest” (Wagner, 2014) and can facilitate adaptation in new selective environments (Gibson and Dworkin, 2004).

Because mutations are random events, it is tempting to picture neutral evolution as a simple random walk (SRW) taking place within neutral networks, and indeed this is how it is usually described, both verbally (Smith, 1970) and in quantitative studies (Huynen et al., 1996). For example, Gavrilets and Gravner address the problem of speciation by considering the percolation of random subgraphs within the sequence hypercube (Gavrilets and Gravner, 1997), assuming that “after a sufficiently long time, the population is equally likely to be at any of the points of the [giant] component” (Gavrilets, 1997). Similarly, Crutchfield and van Nimwegen attempt to link evolutionary dynamics with statistical mechanics via the “maximum entropy” assumption that “[infinite] populations [have] equal probabilities to be in any of the microscopic states consistent with a given [neutral network]” (Crutchfield and van Nimwegen, 2002). These and many other works treat neutral evolution as though its effect were to wash out concentration gradients in genotype space the same way particle diffusion washes out concentration gradients in liquids or gases.

This picture breaks down when many neutral variants co-exist within the same population, *i.e.* when the number of new mutants per generation *M* is much larger than one. It is well known that clonal interference can lead to the loss of beneficial mutations and limits the speed of adaptation in asexual populations (Gerrish and Lenski, 1998; Park and Krug, 2007). Here I show that *neutral interference*—the competition of mutants with equal adaptive value but possibly different robustness through negative selection—has a similar, but perhaps more counter-intuitive, effect on neutral evolution. Instead of allowing a population to explore its neutral network at a faster rate, increasing *M* can confine it to a small, highly connected region within that network. This effect is conceptually and mathematically similar to the Anderson conductor-insulator localization transition in disordered metals (Anderson, 1958) and can be captured with a different kind of random walk model—a “maximal entropy random walk” (Burda et al., 2009).

## Results

### Neutral evolution in a holey landscape

The simplest setting to study neutral evolution is a *holey fitness landscape* (Gavrilets, 1997; van Nimwegen et al., 1999), understood as a set of genotypes with binary fitness *w*: a genotype is either fully functional (*w* = 1), or it is unviable (*w* = 0). Representing possible mutations as edges turns this landscape into a graph. Generically, the subset of functional genotypes in a holey landscape splits in multiple connected components, one of which (the “giant component” *G*) contains the majority of all genotypes. We denote *A* the adjacency matrix of *G* and *d*(*x*) = ∑_*y*∈*G*_ *A*_*xy*_ the neutral degree (number of neutral mutants) of a genotype *x*.

In the classical description of neutral evolution (Kimura, 1983), a functional mutant has a probability 1*/N* to fix and replace the wild type through genetic drift. Under this substitution process the entire population performs a SRW within *G* (with *N* -independent jump rate), whence the concept of populations “diffusing in a neutral network” evoked earlier. However, when the number of new mutants *M* = *µN* ≫ 1, as in e.g. RNA viruses (Drake and Holland, 1999), the selection of mutational robustness becomes more important than genetic drift as a driver of evolution (Schuster and Swetina, 1988; van Nimwegen et al., 1999; Forster et al., 2006; Sanjuán et al., 2007). The dynamics of the distribution of viable genotypes *p*_*t*_ (*x*) is then better described by a replicator-mutator (or “quasi-species”) equation of the form (van Nimwegen et al., 1999)

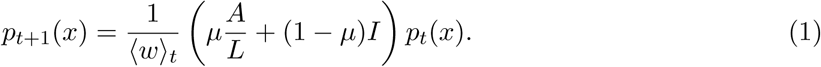

where *L* is the total number of possible mutants for each sequence and the population mean fitness ⟨*w*⟩_*t*_ = *µ*⟨*d*⟩_*t*_*/L* + (1 − *µ*) ensures that *p*_*t*_(*x*) is normalized in *G*. From 1 it is easy to see that the equilibrium distribution (or mutation-selection balance) *Q*(*x*) is an eigenvector of the adjacency matrix of the neutral network *A*; by the Perron-Frobenius theorem *Q*(*x*) is in fact the dominant eigenvector of *A* and the equilibrium mean neutral degree ⟨*d*⟩_∞_ is its spectral radius *ρ*, both of which only depend on the topology of *G*. Since the eigenvector *Q*(*x*) tends to concentrate in regions with high connectivity, these facts establish the “neutral evolution of mutational robustness” (van Nimwegen et al., 1999).

### Equivalence with the maximal entropy random walk

To understand the limits of neutral evolution in interfering populations we must study the full trajectory *p*_*t*_(*x*) and not just its asymptotic equilibrium *Q*(*x*). For this purpose we can use the framework recently developed in (Smerlak, 2019), which maps the non-linear dynamics 1 onto a Markov chain on *G* via the change of variables *q*_*t*_(*x*) ∝ *Q*(*x*)*p*_*t*_(*x*) (SI). This new distribution satisfies the master equation

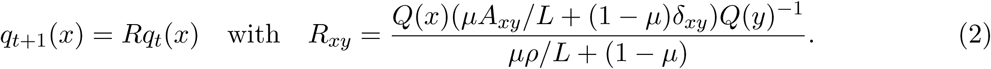

This equation describes a stochastic walk on *G* such that, at each time step, the walker either stays in her current location *x* or jumps to a neighboring node *y* in *G* with probability proportional to *Q*(*y*)*/Q*(*x*). The sequence of actual transitions is then a *maximal entropy random walk* (MERW) on *G*_0_ (SI).

The MERW was introduced as a Markov process with optimal mixing properties by Burda *et al.* (Burda et al., 2009) and has found applications in complex networks theory, image analysis and other fields. In a nutshell, while a simple random walker is “blind” (or “drunk”) and therefore chooses the next node to visit uniformly at random among nearest neighbors, a maximal-entropy random walker is “curious”: her transition probabilities are such that the each new step is as surprising as possible, *i.e.* the MERW maximizes the entropy *rate* of the process. Somehow paradoxically, the blind walker is sure to eventually visit all nodes of a (finite, connected) graph with finite probability, but the curious walker may not: in irregular networks, the equilibrium probability of the maximum-entropy random walker is exponentially suppressed outside a small “localization island” (Burda et al., 2009), as in Fig. S1. In the context of neutral evolution, this implies that narrow neutral ridges are much more difficult to traverse than is usually appreciated— percolation is not enough.

### Revisiting Maynard Smith’s four-letter model

To illustrate this localization effect I reconsidered Maynard Smith’s famous toy model of protein evolution (Smith, 1970). In the set of all possible four-letter words, Maynard Smith used meaning as a binary measure of fitness: any meaningful word is considered functional. He gave the sequence of one-point mutations *σ* = (**word, wore, gore, gone, gene**) as an example of a neutral path, arguing that, unless a large section of genotype space can be traversed through such neutral paths, molecular evolution is impossible. It is easy to come up with other examples of neutral paths; in the following we will focus on *σ*′ = (**opus, onus, anus, ants, arts**).

Using the Wolfram dataset of English “KnownWords” we find that, out of 2405 meaningful four-letter words, 2268 belong to the giant component *G*, including both paths *σ* and *σ*′. However, due to the irregular structure of *G*_0_ with three communities separated by narrow ridges (Fig. 2A), the majority of these words have negligible equilibrium probability: a core of just 420 (resp. 1064) words concentrates 90% (resp. 99%) of the total probability (Fig. S2). In particular, if all the words in Maynard Smith’s path *σ* (except **gene**) belong to the 99%-core, none of the words in *σ*′ do. Note that the words with the largest equilibrium probability are poorly predicted from their neutral degree, as illustrated in Fig. 2B: while **says** and **seed** both have relatively high neutrality (*d* = 21), the former is ten thousand times more frequent than the latter. This highlights that the “neutral evolution of mutational robustness” is not simply the evolutionary advantage of robust genotypes—it is a selection principle which singles out, on a logarithmic scale, a subset of robust (e.g. **bays**) and non-robust (e.g. **whys**) genotypes that are globally well-connected in *G*. These patterns generalize to words of different lengths (Fig. S4 and Table S1).

**Figure 1:**
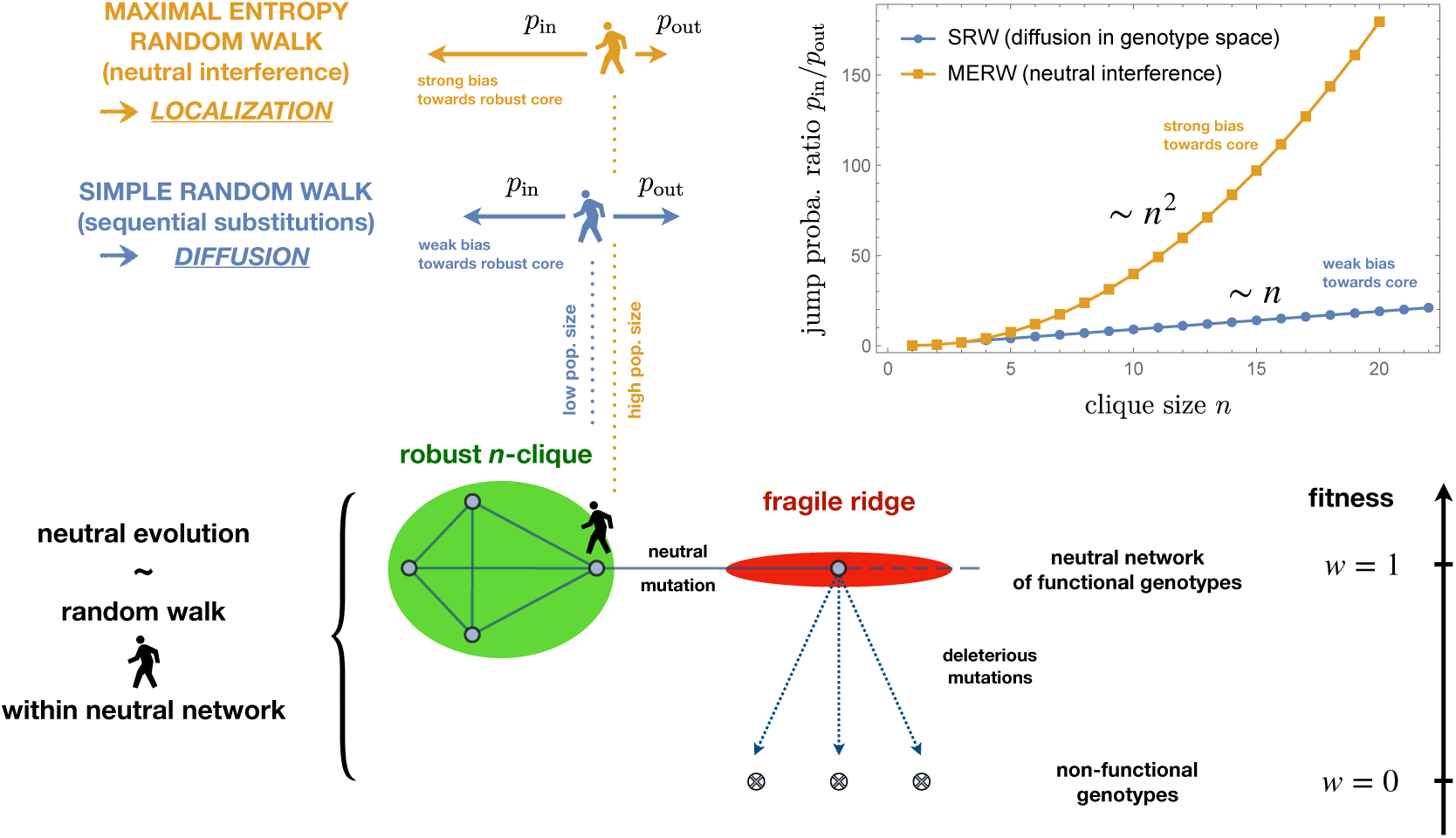
Two Markovian limits of neutral evolution with sharply different dynamics. When mutations are rare and populations mostly homogeneous, neutral evolution amounts to a simple random walk (SRW) along a neutral network *G*. In larger populations, the interference of multiple neutral mutants leads to the selection for mutational robustness, which can be described as a maximal entropy random walk (MERW) on *G*. The qualitative difference between the SRW and the MERW is illustrated here by a simple network configuration where an *n*-clique is connected to a one-dimensional ridge. In that case, the SRW on the edge of the clique with jump back into it with probability *p*_in_ ∼ *np*_out_, while the MERW will jump back with the much higher probability *p*_in_ ∼ *n*^2^*p*_out_. This stronger attraction towards robust cores can induce the localization of populations within *G*, invalidating commonly analogies between neutral evolution and diffusion (or percolation) in sequence space.

**Figure 2:**
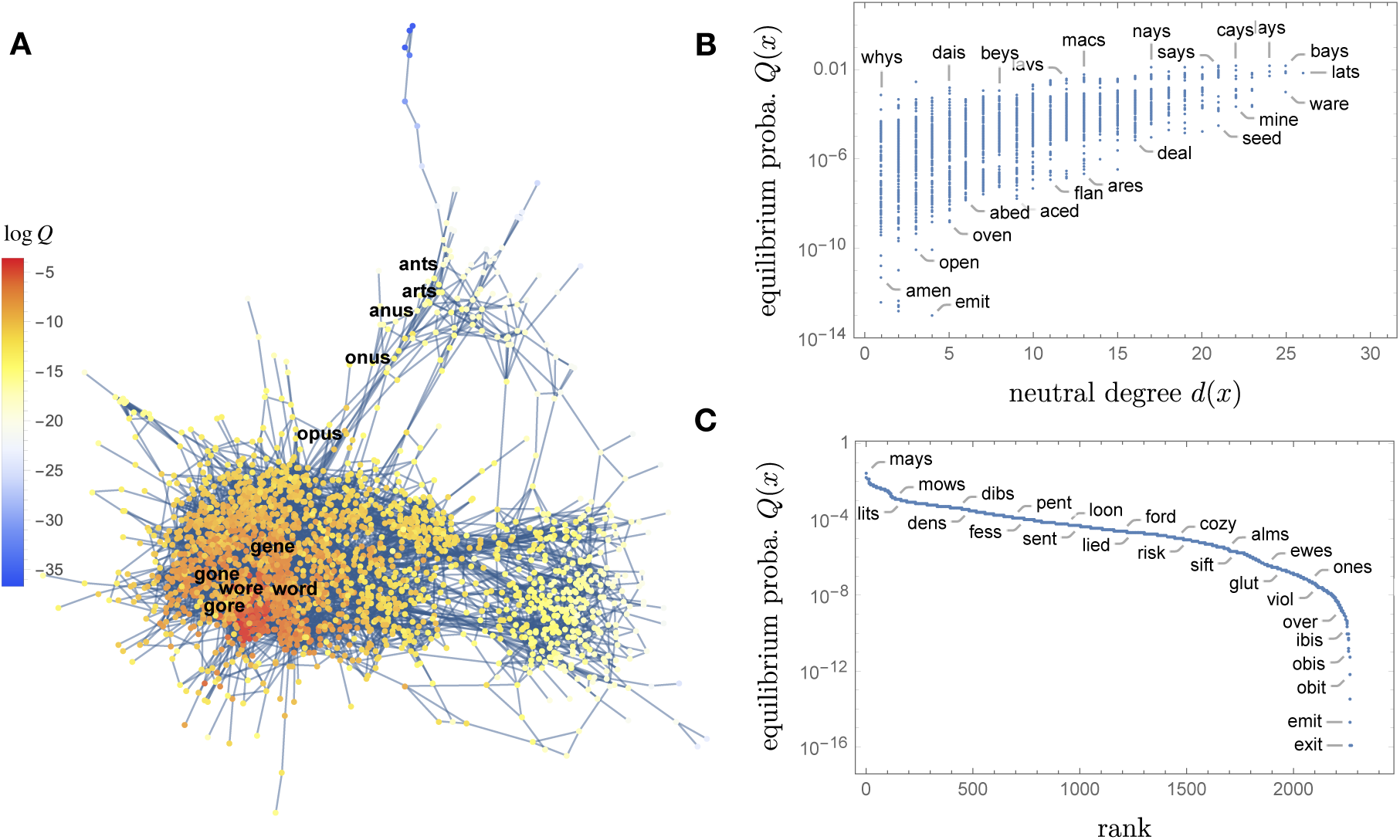
Localization in Maynard Smith’s holey landscape of meaningful four-letter words. A: The giant component with vertices colored by their logarithmic probability at mutation-selection balance, with the neutrals paths *σ* and *σ*′ highlighted in boldface characters. While *σ* is deep in the core and easily evolvable, *σ*′ belongs to a narrow ridge which can hardly be traversed. B: The evolutionary stability of four-letter words, measured by their probability at mutation-selection equilibrium, is correlated with their neutral degree (mutational robustness), but cannot be reliably inferred from it. C: The rank plot of *Q*(*x*) shows the approximately exponential decay of the equilibrium density away from the core typical of localization phenomena.

In multiple runs of a simple evolutionary simulation with a 10% mutation probability per genome per generation, population size *N* = 10^3^ and time horizon of 10^4^ generations, **gene** almost always evolved from **word** (through Maynard Smith’s or, more often, some other path), but—because of the fragility of its intermediate forms—**arts** only rarely evolved from **opus** (Fig. 3A and S3). When it did evolve, **arts** arose in the first few generations of the run, before the population got permanently trapped in the core. Strikingly, smaller (but not too small) populations had a *higher* probability to evolve **arts** at least once within the prescribed time horizon (Fig. 3B). This effect is due to the coupling between individuals through negative selective pressures and would be incomprehensible if we view neutral evolution as diffusion.

**Figure 3:**
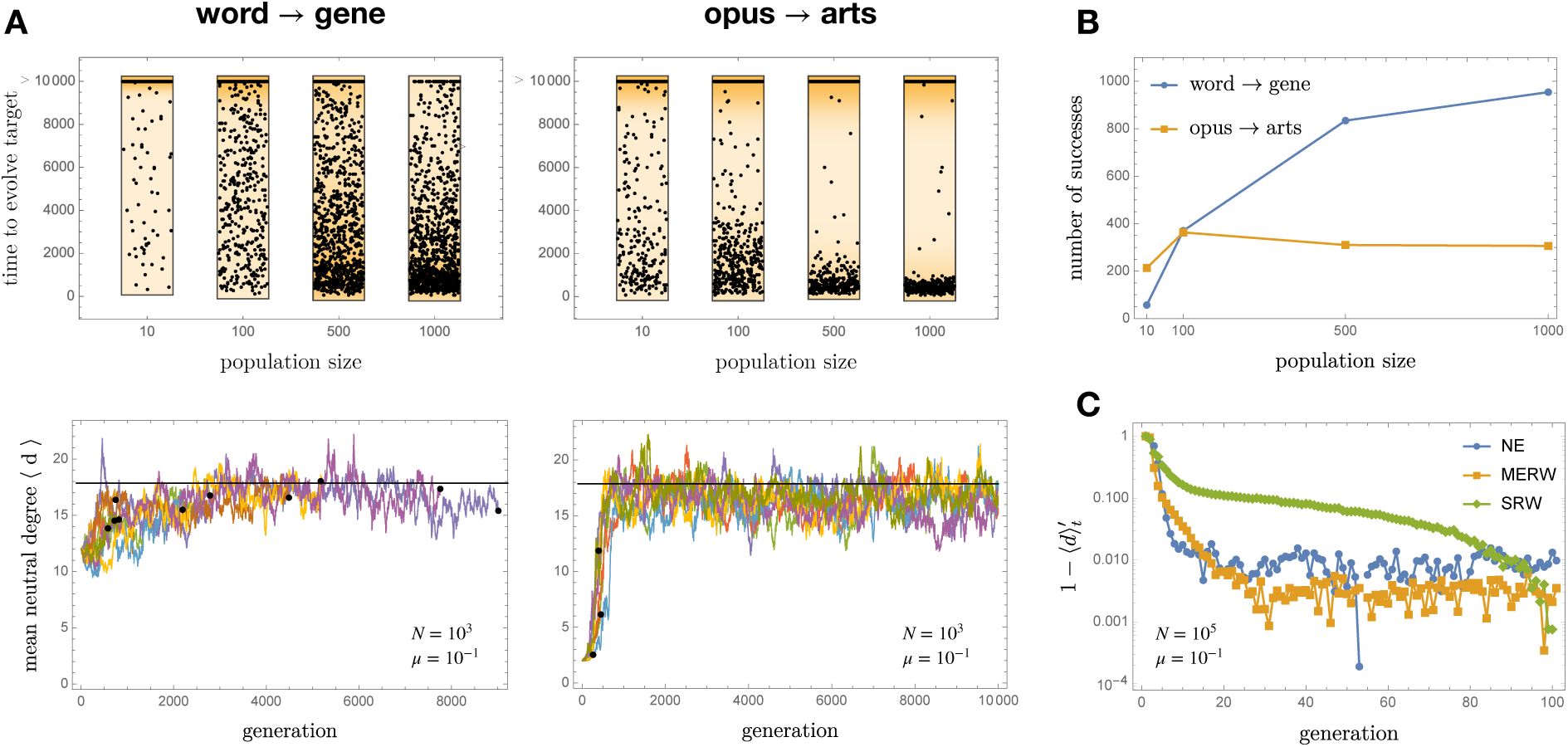
A: The evolution of arts from opus (left column) is a very different challenge than that of gene from word (right column), as is seen by comparing the time to evolve the target word at varying *N* (top row, with a cutoff after 10^4^ generations) and the shape of ⟨*d*⟩_*t*_ trajectories at *N* = 10^3^ (bottom row, with the black horizontal line representing the equilibrium value *ρ* and black dots the endpoint of successful trajectories). While the likelihood of success of word → gene increases with the population size, it does *not* for opus → arts. If a population does not succeed to evolve arts in the first few generations of the run, before it can get trapped in the core, it never will. B: The likelihood to evolve a genotype from another through neutral evolution can depend non-monotonically on the population size. Here there were 10^3^ attempts with *µ* = 0.1, and a population with 100 individuals was more likely to evolve opus → arts at least once than one with 1000 individuals. C: The convergence to equilibrium, here measured by the rescaled mean degree 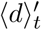, is much faster in a neutrally evolving population (NE) than in diffusing one (SRW) of the same size; it is however consistent with the maximal mixing property of the MERW.

### RNA secondary structures

As a second illustrative example I reconsidered the neutral network of RNA secondary structures studied in (van Nimwegen et al., 1999). The secondary structure of an RNA molecule is the pattern of pairings between complementary bases along its sequence, and can be represented with brackets (paired bases) and dots (unpaired bases). Here we consider sequences of length *ℓ* = 18 with minimum-free-energy secondary structure “((((((….)))..)))” and only purine-pyrimidine base pairs. With the RNA Vienna folding algorithm v2.4.8 (Hofacker, 2003), the giant component has size |*G*_0_| = 17557, but its mutation-selection equilibrium distribution *Q*(*x*) varies over 7 orders of magnitude between core and periphery due to narrow ridges within *G* (Fig. 4); such ridges are also seen in the experimental assay of a small protein neutral network (Podgornaia and Laub, 2015), and may be a generic feature of biological genotype-to-phenotype mappings (Aguirre et al., 2011). Thus, in molecular evolution as well as in Maynard Smith’s toy model, neutral evolution may not efficiently sample neutral networks. Statements to the effect that “diffusion enables the search of vast areas in genotype space” (Huynen et al., 1996) must therefore be qualified accordingly.

**Figure 4:**
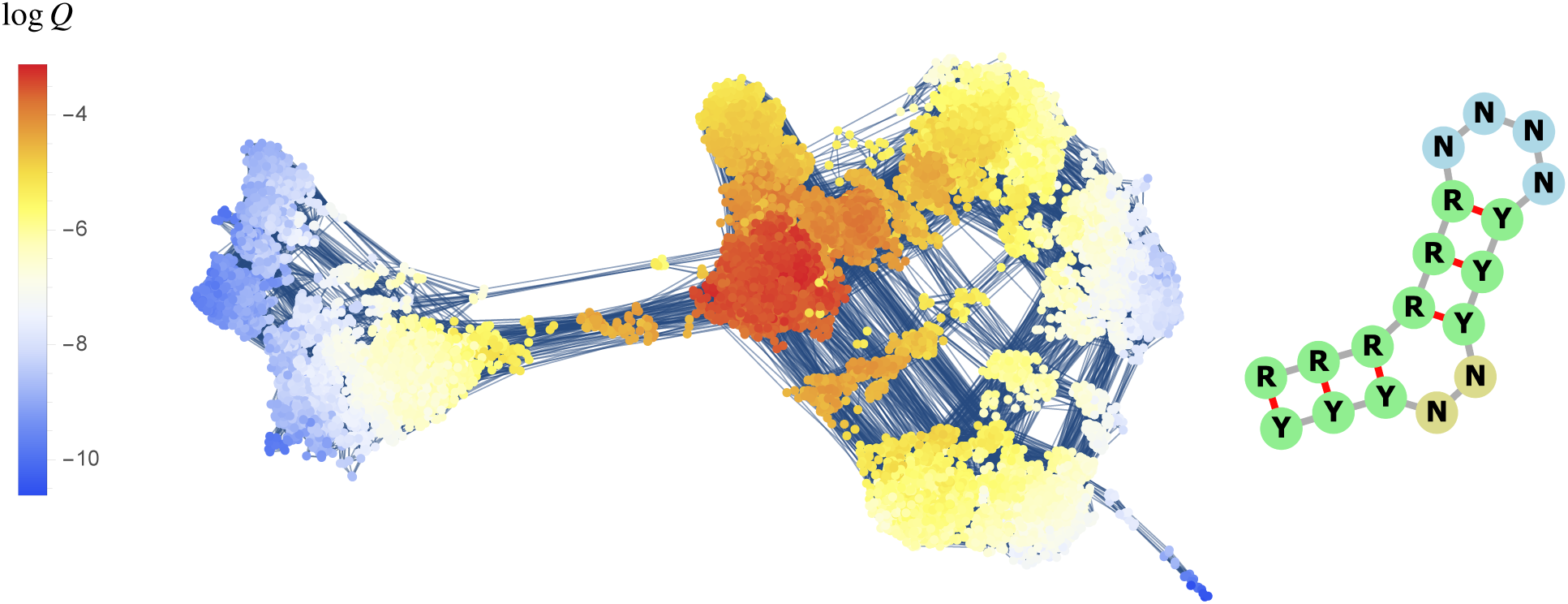
The neutral network of the RNA secondary structure “((((((….)))..)))” with only purine (R) - pyrimidine (Y) base pairs (van Nimwegen et al., 1999), as predicted by the RNA Vienna algorithm v2.4.8. Its giant component consists of multiple communities separated by narrow ridges which induce the localization of the mutation-selection equilibrium *Q* and limits non-equilibrium dynamics accordingly. For this reason the notion that “diffusion enables the search of vast areas in genotype space” (Huynen et al., 1996) must be understood as holding within communities but not between them.

### Robustness and evolvability

The association of NE with the MERW rather than the SRW has notable implications for the classical issues of robustness and evolvability (Masel and Trotter, 2010). In the literature, the potential of (e.g. RNA) sequences to generate new phenotypes—their evolvability—has been related to the (linear or logarithmic) size |*G*_0_| of their neutral network (Jörg et al., 2008), aka their versatility. The rationale for this hypothesis is that a large neutral network potentially communicates with many other neutral networks (other phenotypes) through “portal” sequences interfacing between them. But, just like the randomness of an information source should be quantified by its entropy and not merely its alphabet size (Shannon, 1948), the reach of neutral evolution should be quantified by its ability to generate new sequences, not by the number of all possible sequences. Information theory provides the correct language for this: in the *M* ≫ 1 regime, the versatility of a phenotype should be measured by the entropy *H*(*Q*) = − ∑_*x*∈*G*_ *Q*(*x*) log *Q*(*x*) of its mutation-selection equilibrium, or equivalently by the effective size |*G*|_eff_ = 2^*H*(*Q*)^ of the giant component. In both cases considered above this quantity is much smaller than the naive value |*G*|: with four-letter words we have |*G*|_eff_ = 346.6 ≪ 2268, and for the RNA secondary structure |*G*|_eff_ = 3971.8 ≪ 17557.

## Conclusion

I have described the localization of populations within neutral networks induced by the competition for mutational robustness among neutral mutants. This “neutral interference” is interference in the two senses of the word: in the biological sense, because it involves the competition of clonal subpopulations, an effect usually referred to as clonal interference; and in the physical sense, because localization is an interference phenomenon normally encountered in (classical or quantum) wave mechanics. The link between these two seemingly different process—the evolution of molecular populations and the destructive interference of waves—is provided by the MERW, a Markov chain whose local transition probabilities depend on the global structure of the underlying graph.

These observations reveal sharper constraints on the navigability of neutral networks than previously appreciated: not only does evolution favor high mutational robustness and low genetic loads, it also positively refuses to engage in tightrope walking along narrow neutral ridges. This effect manifests itself in the surprising non-monotonic dependence of discovery rates of certain target genotypes on population size and implies that hopes to “infer the complete structure of the neutral network from accurate measurements of the transient population dynamics” (van Nimwegen et al., 1999) are unfounded. In this way we can only hope to infer the structure of the robust cores of neutral networks. In molecular fitness landscapes, walking a narrow ridge is not *a priori* easier than crossing a fitness valley.

## Acknowledgements

I thank the members of the Structure of Evolution group at MPI MiS, and especially A. Zadorin and I. H. Hatton, for useful critical comments. Funding for this work was provided by the Alexander von Humboldt Foundation in the framework of the Sofja Kovalevskaja Award endowed by the German Federal Ministry of Education and Research.

## Supplementary Information

### Selection-mutation dynamics as a Markov process

In (Smerlak, 2019) I showed that continuous-time replicator-mutator (or quasispecies) equations can be understood in terms of a derived Markov process, in which the logarithm of the selection-mutation equilibrium plays the tole of an effective potential. With discrete generations, this scheme can be reformulated as follows. Given a discrete space *X*, consider a sequence of probability distributions *p*_*t*_ : *X* → ℝ evolving under the dynamics

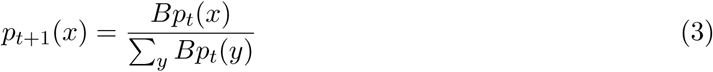

with *B* any irreducible non-negative matrix. (Here I identify a function on *X*with the vector of its values.) In replicator-mutator systems we have *B* = *MW* with *W* a diagonal matrix of Wrightian fitnesses and *M* a stochastic matrix of mutation probabilities. The process 3 is not Markovian due to the global interactions introduced by the normalization factor; for this reason it is difficult to interpret it in terms of “evolutionary trajectories”.

This can be remedied by means of the change of variable *q*_*t*_(*x*) ∝ *S*(*x*)*p*_*t*_(*x*), where *S* is the (left) eigenvector of *B* with largest eigenvalue; by the Perron-Frobenius theorem this vector is positive and its eigenvalue is the spectral radius *σ* of *S*. Via this transformation—which only depends on *t* through a global constant—we obtain a Markovian representation of the original dynamical problem with master equation

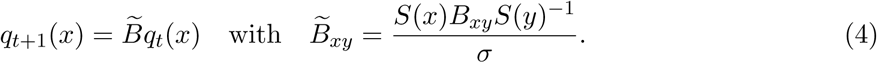

For this derived process the function *U* = −2 log *S* plays the role of a potential; its analysis reveals the metastable states and preferred trajectories of the original (non-linear) process (Smerlak, 2019). The equilibrium distribution for 4 is *q*_∞_(*x*) ∝ *S*^2^(*x*), and the original distribution can be reconstructed from *q*_*t*_(*x*) with *p*_*t*_(*x*) ∝ *S*(*x*)^−1^*q*_*t*_(*x*). When *B* happens to be the adjacency matrix *A* of a connected graph *G*, as in neutral evolution, the process 4 coincides with the MERW on *G*.

### The maximal entropy random walk

The SRW on a connected graph *G* with adjacency matrix *A* and degree matrix *D* = diag(*d*(*x*))_*x*∈*G*_ is the Markov chain with transition matrix

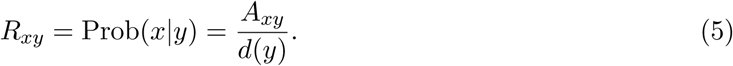

By contrast, the MERW was defined in (Burda et al., 2009) as the Markov chain with transition matrix

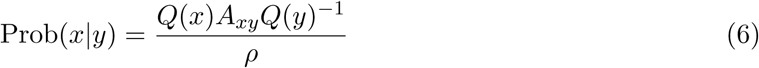

where *Q* is the dominant eigenvector of *A* with eigenvalue *ρ* (equal to the spectral radius of *A*). This Markov chain maximizes the entropy rate among ergodic stationary processes on *G* by assigning equal probability to all paths connecting two given nodes. Rather paradoxically, this property of “maximal mixing” also leads to dynamical localization when the graph *G* is irregular, as in Fig. S1.

### Attraction to a clique in the SRW and MERW

The qualitative difference between the SRW and the MERW can be understood by considering the probabilities *p*_in,out_ for a walker to jump into (resp. out of) a fully connected subgraph with *n* nodes (an *n*-clique) along a ridge, as in Fig. 1. Since there are *n*−1 edges going in and 1 edge going out, for the SRW these probabilities are simply 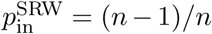 and 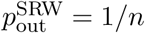, hence 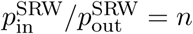.

To evaluate the same ratio for the MERW, we must compute the dominant eigenvector *Q* of the adjacency matrix *A* for the complete graph over *n* nodes {1, …, *n*} with one extra node, labelled 0, attached to vertex 1. By symmetry this eigenvector *Q* = (*q*_0_, *q*_1_, … *q*_*n*_) can be chosen such that *q*_2_ = … = *q*_*n*_ = 1. Writing *AQ* = *ρQ* then gives a system of two quadratic equations for (*q*_0_, *q*_1_), from which we then obtain 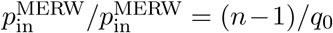. Because *q*_0_ ∼ 1*/n* when *n* ≫ 1, this gives 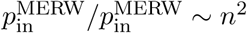 as claimed in Fig. 1.

### Evolutionary simulations

The simulations displayed in Fig. 2 and S3 were performed with the following algorithm, which takes as input a connected graph *G*, an integer *δ* ≥ max_*x*∈*G*_ *d* (*x*) representing the total number of possible (neutral or lethal) mutants a genotype can have, a population size *N*, a mutation probability *µ*, a time horizon *T*, an initial genotype *x*_0_ ∈ *G* and a target genotype *x*_1_ ∈ *G*. Here we denote for each *x* ∈ *G ν* (*x*) the set of its *d* (*x*) neighbors in *G*, and let 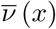 be the set obtained by adjoining *δ* − *d* (*x*) copies of the symbol † (meaning “dead”) to *ν* (*x*).

First, generate an initial population *P*_0_ consisting of *N* copies of *x*_0_. Next, for each generation *t*, perform the two following steps:

- **mutation**: draw a number *n* ∼ Binom(*N, µ*) and replace *n* randomly chosen individuals *x* in *P*_*t*_ by random samples from the corresponding 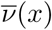; these samples together with the elements of *P*_*t*_ not chosen for mutation form the mutated population 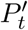
- **selection**: sample with replacement *N* elements from 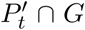 and collect them in a new population *P*_*t*+1_

The algorithm terminates whenever *x*_1_ ∈ *P*_*t*_ (the target genotype is found) or *t* = *T* (the time horizon is reached).

## Supplementary Figures

**Figure S1:**
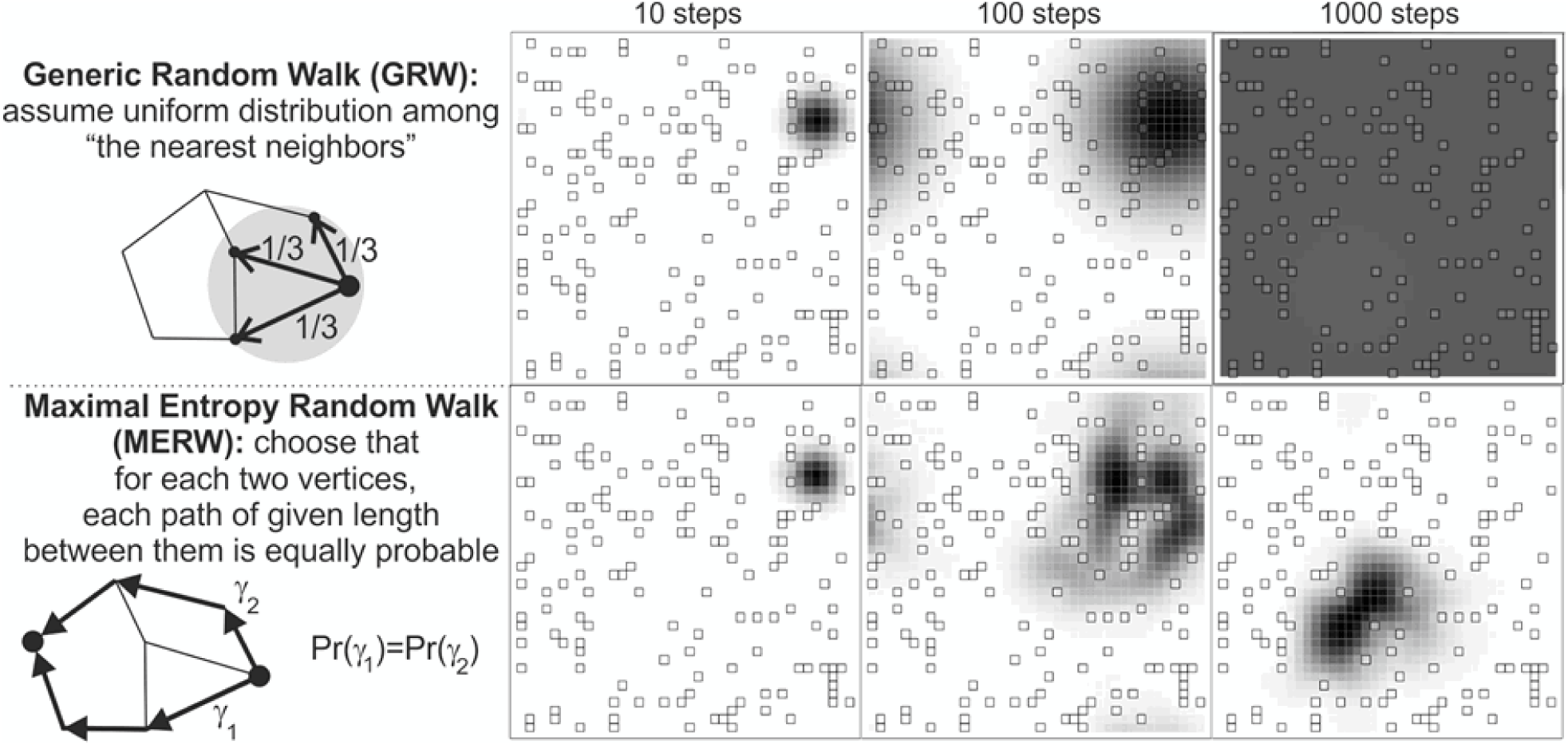
Illustration of the difference between the simple (or generic) random walk (top) and the maximal entropy random walk (bottom) in a two-dimensional lattice with defects. The SRW choose elementary steps uniformly at random among nearest neighbors and eventually fills the lattice uniformly (top); the MERW chooses paths between fixed nodes uniformly at random and ends up trapped in a defect-poor “localization island” (bottom). Illustration by Jarek Duda (CC BY-SA 4.0).

**Figure S2:**
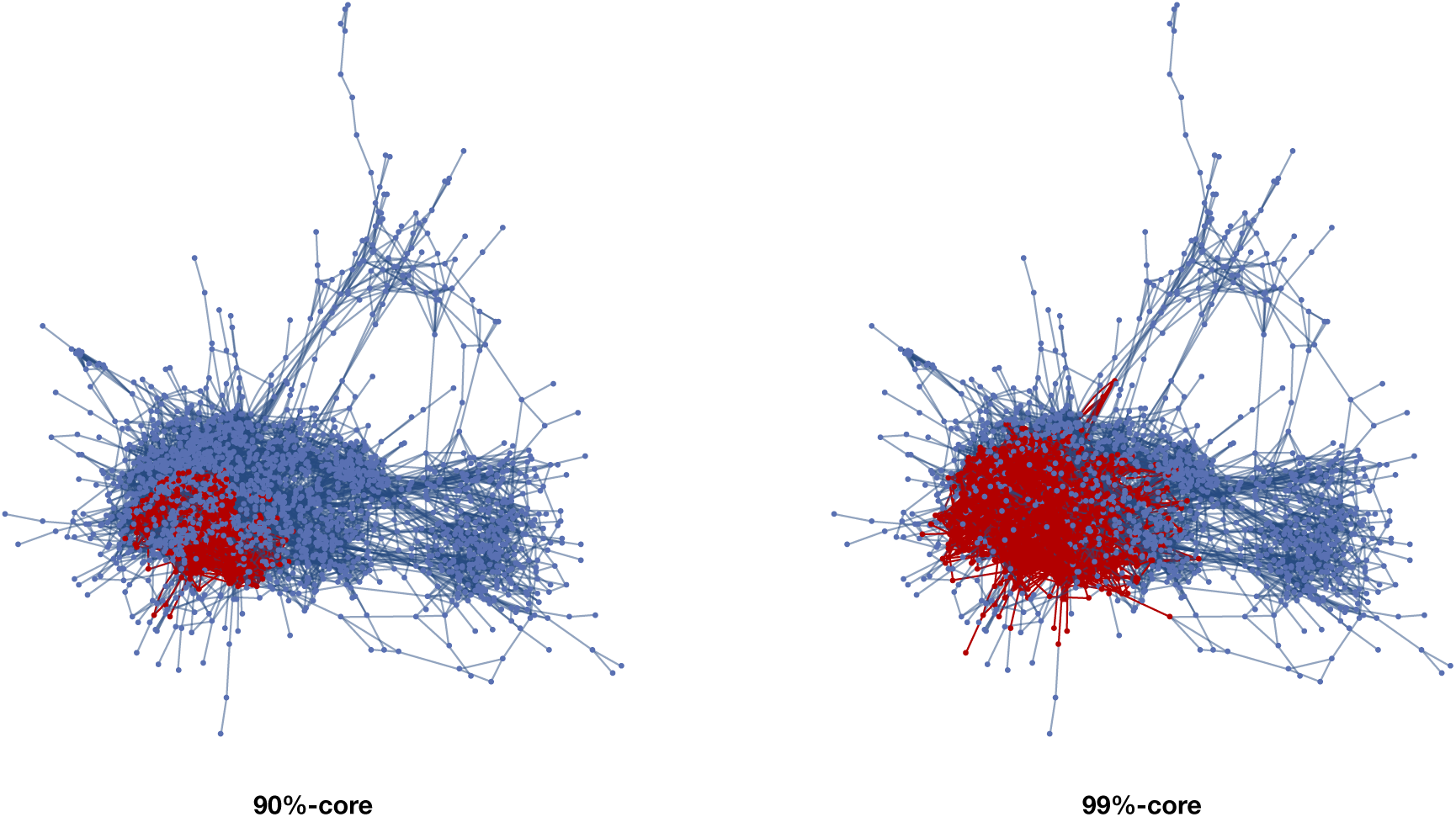
The 90%- and 99%-cores within the giant component of meaningful four-letter English words, with size 420 and 1024 respectively (out of 2268 words in the giant component and 2405 in total). These cores correspond to regions with higher mutational robustness (van Nimwegen et al., 1999).

**Figure S3:**
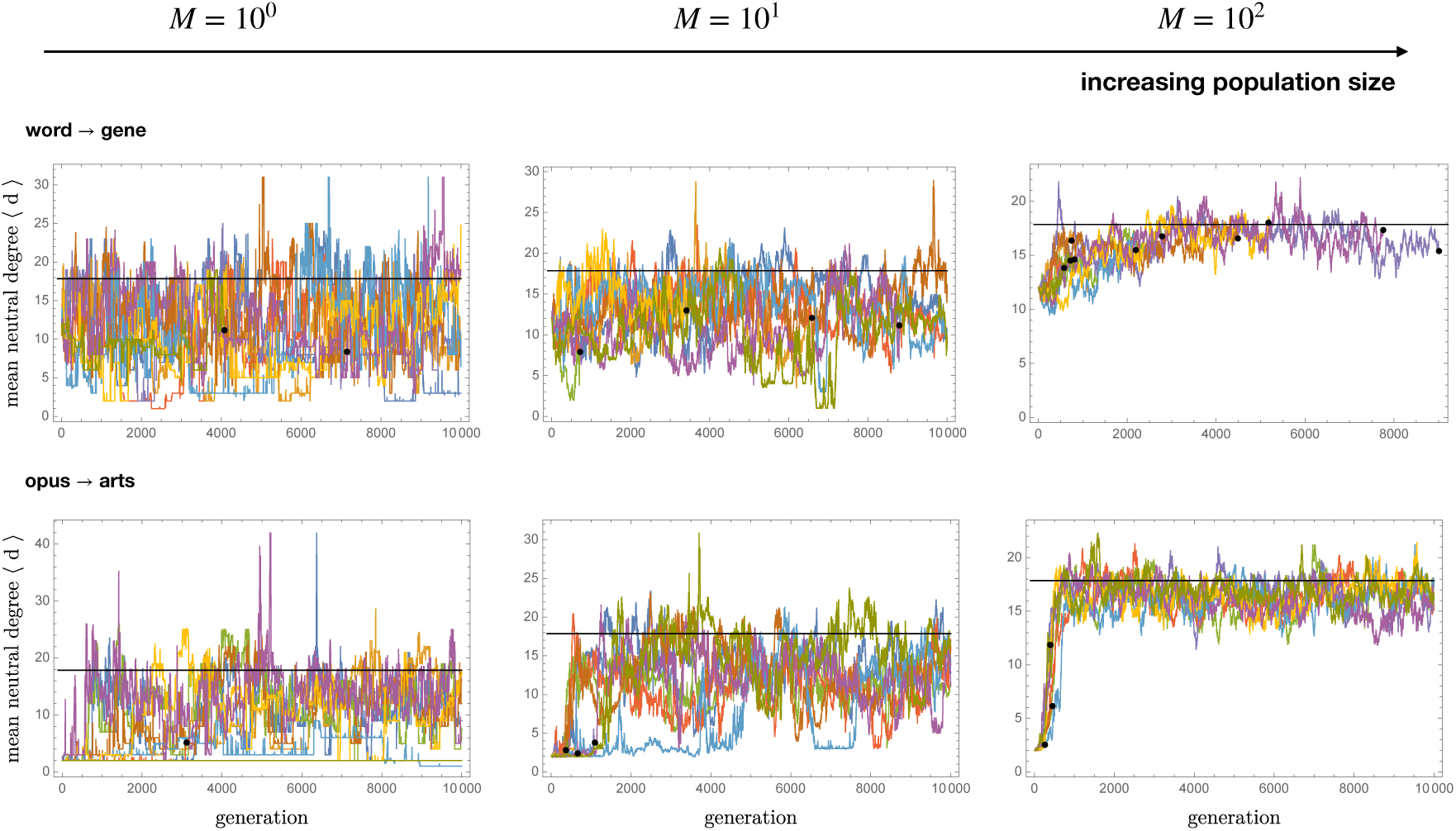
Sample evolutionary trajectories for the **word** → **gene** (top) and **opus** → **arts** (bottom) problems, with success shown as a black dot. When just a few mutants co-exist in the population (*M* = 1), neutral evolution is insensitive to mutational robustness and behaves as a SRW, blindly exploring the whole neutral network. As *M* increases, the attraction towards the core of the giant component (where ⟨*d*⟩ = *ρ*, indicated by the black horizontal line) becomes stronger and the likelihood to walk a tightrope such as *σ′* decreases accordingly.

**Figure S4:**
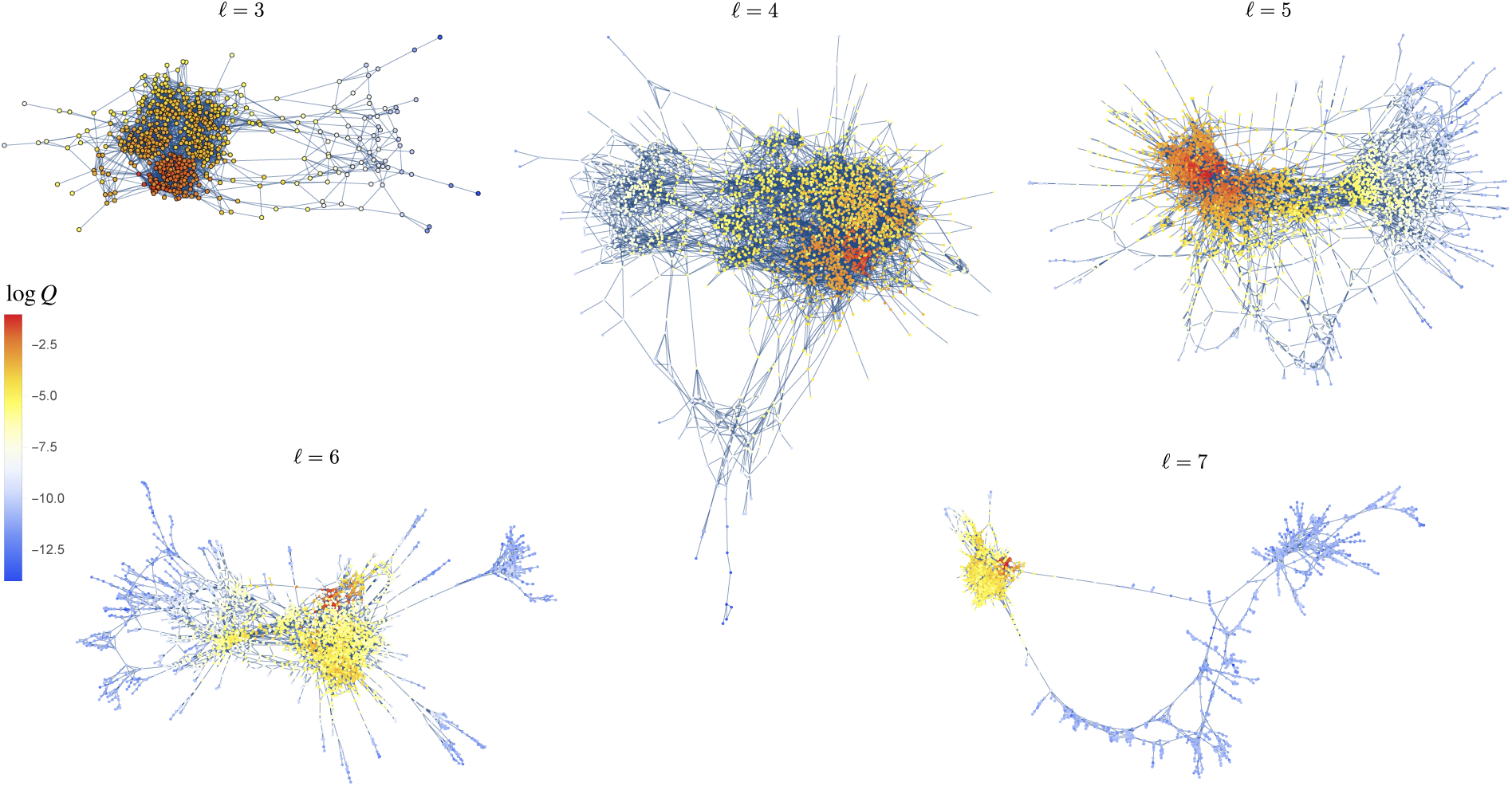
Giant components in the holey landscapes of meaningful English words of different lengths *ℓ*, with words *x* colored by their logarithmic mutation-selection equilibrium probability *Q*(*x*). In all cases the latter displays exponentially localization in a robust core.

## Supplementary Table

**Table S1:**
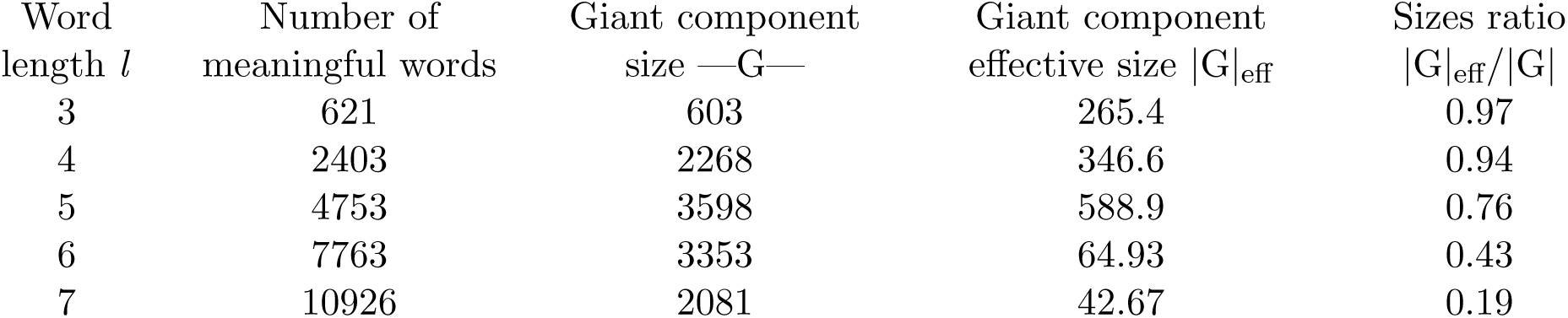
Holey landscapes of meaningful English words of different lengths, see Fig. S4.

